# DNA methylation in transposable elements disrupts the connection between three-dimensional chromatin organization and gene expression upon rice genome duplication

**DOI:** 10.1101/2021.12.15.472849

**Authors:** Zhenfei Sun, Yunlong Wang, Zhaojian Song, Hui Zhang, Min Ma, Pan Wang, Yaping Fang, Detian Cai, Guoliang Li, Yuda Fang

**Affiliations:** Joint Center for Single Cell Biology, School of Agriculture and Biology, Shanghai Jiao Tong University, Shanghai 200240, China; National Key Laboratory of Crop Genetic Improvement, Hubei Key Laboratory of Agricultural Bioinformatics, College of Informatics, Huazhong Agricultural University, Wuhan, 430070, China; School of Life Science, Hubei University, Wuhan 430062, China; CAS Center for Excellence in Molecular Plant Sciences, Institute of Plant Physiology and Ecology, Chinese Academy of Sciences, Shanghai 200032, China

**Keywords:** Polyploidy rice, Hi-C, TAD, transposable elements, DNA methylation

## Abstract

Polyploidy serves as a major force in plant evolution and domestication of cultivated crops. However, the relationship and underlying mechanism between three-dimensional (3D) chromatin organization and gene expression upon rice genome duplication is largely unknown. Here we compared the 3D chromatin structures between diploid (2C) and autotetraploid (4C) rice by high-throughput chromosome conformation capture analysis, and found that 4C rice presents weakened intra-chromosomal interactions compared to its 2C progenitor. Moreover, we found that changes of 3D chromatin organizations including chromatin compartments, topologically associating domain (TAD) and loops uncouple from gene expression. Moreover, DNA methylations in the regulatory sequences of genes in compartment A/B switched regions and TAD boundaries are not related to their expressions. Importantly, in contrast to that there was no significant difference of methylation levels in TEs in promoters of differentially expressed genes (DEGs) and non-DEGs between 2C and 4C rice, we found that the hypermethylated transposable elements across genes in compartment A/B switched regions and TAD boundaries suppress the expression of these genes. We propose that the rice genome doubling might modulate TE methylation which results in the disconnection between the alteration of 3D chromatin structure and gene expression.

## Introduction

Polyploidy plays an important role in the formation of plant new species, evolution and breeding (Marcussen et al., 2014; Soltis et al., 2015). Polyploidy plant is often accompanied by powerful biological potentials, improved environmental adaptation, elevated biomass and yield (Chao et al., 2013; Wu et al., 2014a). It was known that autotetraploid rice shows larger kernels, higher protein content, better amino acid composition and higher 1,000-grain weight than its diploid counterpart (Wu et al., 2014b). Autotetraploid *Arabidopsis* exhibits obvious phenotypes at both vegetative and reproductive stages, including large leaves, increased whole plant size, late flowering and large seeds (Zhang et al., 2019). Autotetraploid birch is superior in volume, leaf, breast-height diameter, fruit and stoma, and inferior in height compared to diploid birch (Mu et al., 2012). At the cellular and molecular levels, polyploidization often leads to changed chromatin structures and gene expression (Chen and Ni, 2006; Zhang et al., 2019; Concia et al., 2020).

Chromatin is organized in a highly ordered three-dimensional (3D) architecture instead of as a linear nucleotide sequence of the genome (Meaburn and Misteli, 2007; Lieberman-Aiden et al., 2009). The 3D genome is packed in the nucleus in a hierarchical pattern. Chromosome territory (CT) at the several megabase-scale is a higher level of chromatin domain (Cremer et al., 2006; Cremer and Cremer, 2010; Gibcus and Dekker, 2013). Chromatin in CT is divided into two types of compartments, A and B. Compartment A is associated with open chromatin and active transcription, and compartment B with closed chromatin and inactive transcription (Lieberman-Aiden et al., 2009). Topologically associating domain (TAD) is a predominant structural feature in most organisms (Wang et al., 2015). TADs often represent functional domains, as a given TAD contains the regulatory elements for the genes inside the same TAD (de Laat and Duboule, 2013). Therefore, the integrity of the TAD structure is necessary for gene regulation (Ibn-Salem et al., 2014; Lupianez et al., 2015). The location of TAD boundary is strongly related to the local gene expression, epigenetic landscape and the binding of various insulator proteins (Dixon et al., 2012; Filippova et al., 2014). Chromatin loops which appear at 10 kb to 1 Mb (Fraser, 2006; Phillips and Corces, 2009) function in transcription, replication and recombination (Mukherjee and Mukherjea, 1988).

It has been known that the alterations of chromatin structures are coupled with the changes of gene expressions in some biological progresses (Ouyang et al., 2020). For example, in *Arabidopsis*, the switches of compartment A/B lead to the change of transcription during the genome doubling (Zhang et al., 2019). In rice, higher-order chromatin architecture is correlated with transcriptional regulation under heat stress (Liang et al., 2021). In cotton, the changes of TADs affect the transcriptional activities of abundant genes in tetraploid compared to diploid cotton (Wang et al., 2018). While other reports have also shown that three-dimensional structural changes are unrelated with gene expression. For example, the uncoupling relationship between genome topology and gene expression was observed in highly rearranged chromosomes (balancers) spanning ~75% of *Drosophila* genome (Ghavi-Helm et al., 2019). Most TAD disruptions do not result in marked changes of gene expression in human cancer (Akdemir et al., 2020). Recently, chromatin structure and the regulation of gene expression were found to be independent during development of *Drosophila* (Espinola et al., 2021; Ing-Simmons et al., 2021). However, the mechanism of both correlation and uncorrelation between 3D chromatin structure and gene expression has not been revealed.

Polyploidy events trigger a large number of epigenetic and transcriptional changes in the replicated or merged genome (Seoighe and Gehring, 2004; Paun et al., 2011; Roulin et al., 2013; Becak, 2014; Diez et al., 2014). In addition to the 3D genomic organization, another important epigenetic factor is DNA methylation at cytosine residues which is associated with gene transcription by affecting the binding of chromatin proteins including transcription factors (TFs) to DNA (Moore et al., 2013). The precise regulation of DNA methylation is essential for plant and animal developments (Luo et al., 2013; Smith and Meissner, 2013; Kawashima and Berger, 2014). In plants, in addition to CG context, DNA methylation also occurs in sequence contexts of CHG and CHH with which siRNAs are mainly associated (Feng et al., 2010; Zemach et al., 2010; Feng and Jacobsen, 2011). The majority of DNA methylation is found in transposable elements (TEs) with CG, CHG and CHH contexts to suppress the activities of TEs. Substantial methylation is found in the bodies of active genes, in which generally occurs in the CG context (Law and Jacobsen, 2010). DNA methylation in regulatory sequences, such as promoters and enhancers, often leads to gene silencing (Jones, 2012; Schubeler, 2015; Zhang et al., 2018a).

In this study, we found that the changes of 3D chromatin structures are not related to the transcriptional changes when diploid (2C) rice is duplicated to autotetraploid (4C) rice. In addition, DNA methylation in the regulatory regions of genes in the compartment A/B switched regions or TAD boundaries are not important for their differential regulations of transcription between 2C and 4C rice. By comparing the methylations of TEs adjacent to genes in the compartment A/B switched regions or TAD boundaries and differentially expressed genes (DEGs) between 2C and 4C rice, we revealed that the elevated methylation in TEs adjacent to genes in the compartment A/B switched regions or TAD boundaries suppresses the transcription of these genes upon rice genome duplication.

## Results

### The changes of phenotypes and gene expression upon rice whole-genome duplication

The diploid (2×9311, 2C) and autotetraploid rice (4×9311, 4C) were confirmed by flow cytometry (Supplemental Figure S1). Compared to 2C rice, 4C rice seedlings show no obvious changes of plant height, flag leaf width, and plant weight (Figure 1A-D) with decreased tillering number and increased flag leaf length, 1,000-grain weight, number of effective panicles per plant, grain length, and grain width (Figure 1E-J), similar to the phenotypes of autotetraploid rice *Oryza sativa ssp. indica* cv. *Aijiaonante* previously reported (Zhang et al., 2015). In addition, the nuclei in leaf and root cells of 4C rice are larger than those of 2C rice (Figure 1K-N).

**Figure 1.**
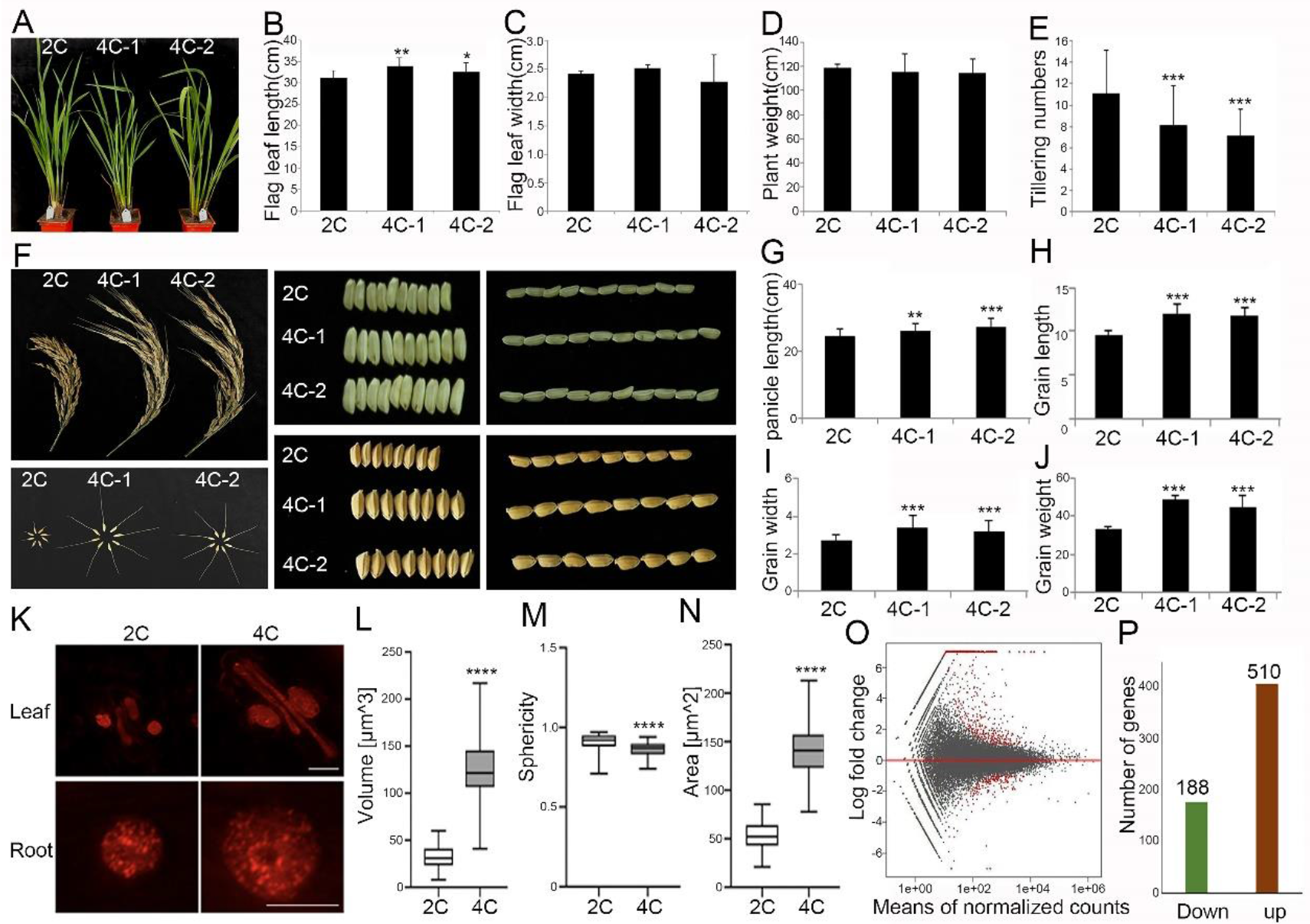
Comparisons of morphology and transcriptome between diploid (2×9311) and autotetraploid rice (4×9311). **A**, Seedling morphology of diploid and autotetraploid rice. **B-E**, Flag leaf length (B), flag leaf width (C), plant weight (D), and tillering numbers (E) of diploid and autotetraploid rice seedlings. Error bars represent means ± SEM (standard error of mean, n = 3 biological replicates). Statistical significance was analyzed by *t*-test, \*\*\**p* < 0.001. **F**, Panicle and grain morphology of diploid and autotetraploid rice. **G-J**, Panicle length (G), grain length (H), grain width (I), and grain weight (J) of diploid and autotetraploid rice plants. Error bars represent means ± SEM (standard error of mean, n = 3 biological replicates). Statistical significance was analyzed by *t*-test, \*\*\**p* < 0.001. **K**, Nuclei of guard cells and root cells of diploid and autotetraploid rice stained by DAPI. **L-N**, Volume (L), sphericity (M), and area (N) of diploid and autotetraploid rice nuclei. Error bars represent means ± SEM (standard error of mean, n = 3 biological replicates). Statistical significance was analyzed by *t*-test, \*\*\**p* < 0.001. **O**, MA plot for statistical significance against gene fold change between diploid and autotetraploid rice. Each gene was marked as a dot. Red dots above 0 represent up-regulated genes, red dots below 0 represent down-regulated genes and black dots represent the other genes. **P**, Numbers of up- and down-regulated genes (|log2fold change| > 1) between diploid and autotetraploid rice seedlings. The data from three biological replicates were combined.

To evaluate the impact of rice genome doubling on transcription, we compared the transcriptomes of the above ground parts between 10 day-old 2C and 4C seedlings by RNA-sequencing. The results indicate that the predominant part of genes is not changed obviously. Among 698 genes significantly regulated, 510 genes are up-regulated and 188 are down-regulated (Figure 1O-P; Supplemental Data Set S1). Gene ontology (GO) analysis indicate that these DEGs associate with a variety of biosynthetic and metabolic processes (Supplemental Figure S2; Supplemental Data Set S2).

### Intra-chromosomal interactions are weakened in autotetraploid rice

To test whether chromatin organization is rearranged after rice genome duplication or not, we performed Hi-C experiments to map the chromatin interactions. More than 843 million and 842 million raw Hi-C reads from 2C and 4C rice were obtained (Supplemental Data Set S3), respectively, with a high reproducibility between the two biological replicates of 2C or 4C rice (Supplemental Figure S3A and 3B). We then calculated relative interaction difference between 2C and 4C rice. The results showed that 4C rice shows slightly increased inter-chromosomal interactions and dampened intra-chromosomal interactions (Figure 2A–2D). We also observed decreased inter- or intra-chromosome arm interactions in most of chromosomes in 4C rice compared to 2C rice (Figure 2E and 2F).

**Figure 2.**
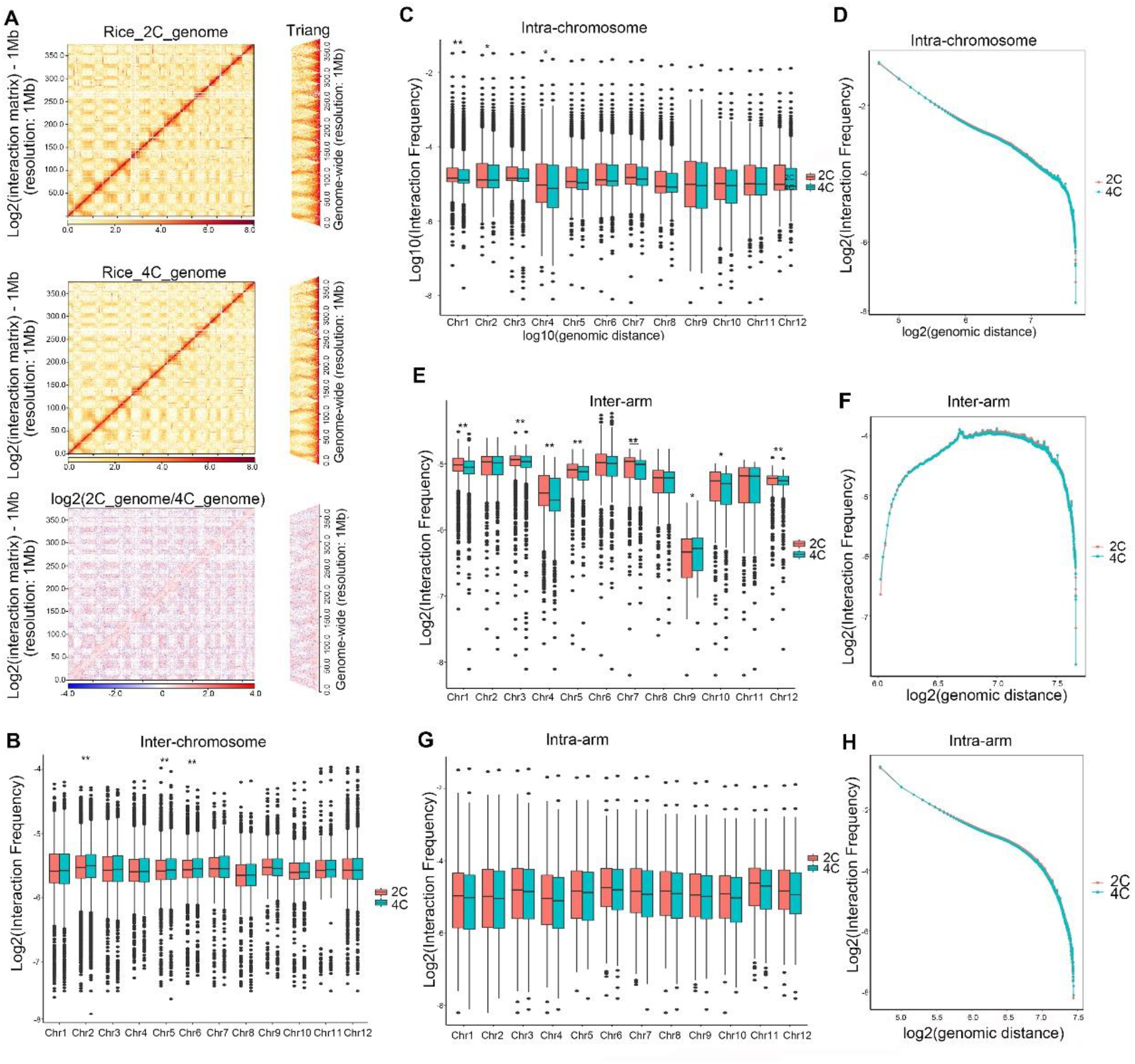
Rice genome doubling weakens the chromatin interactions. **A**, Chromatin interaction heatmaps of 2C and 4C rice, and differential chromatin interaction heatmap between 2C and 4C rice at a 1M resolution. Chromosomes stacked from bottom left to up right were chr1, chr2, chr3, chr4 chr5. **B**, Boxplots showing inter-chromosome interaction frequencies among all chromosome pairs. **C**, Boxplots showing intra-chromosome interaction frequencies between 2C and 4C rice. **D**, Interaction decay exponents of intra-chromosome interactions. **E**, Boxplots showing inter-arm interaction frequencies between 2C and 4C rice. Inter-arm interactions are the interactions with both sides inside one chromosome, but from different arms of the same chromosome. **F**, Interaction decay exponents of inter-arm interactions. **G**, Boxplots showing intra-arm interaction frequencies between 2C and 4C rice. **H**, Interaction decay exponents of intra-chromosome arm. (\*\*\**p* < 0.001, \*\**p* < 0.01, \**p* < 0.05, NS *p*>0.05. The *p* values were tested by Wilcoxon–Mann–Whitney test).

To quantitatively assess the chromatin contacts, we calculated interaction decay exponents (IDEs), which characterize chromatin packing as the slopes of a linear fit of average interaction intensities detected at a given range of genomic distances in the logarithm scale (Grob et al., 2014). The results displayed that IDEs of intra-chromosomes (Figure 2D; Supplemental Figure S4), inter-chromosome arms (Figure 2G), intra-chromosome arms (Figure 2H) in 4C rice are slightly lower than those in 2C rice.

### The switches between chromatin compartment A and B upon rice genome doubling are not related to transcriptional regulation

To know whether the chromatin compartment change is associated with the alteration of gene expression during rice genome duplication, we used first principal component of the Pearson's matrix in Hi-C data and gene expression to define the active (A) and inactive (B) chromatin compartments in 2C and 4C rice. Compartments A and B were compared through the first principal component at a 50 kb resolution between 2C and 4C rice. The results indicate that 47.31% and 50.3% of the genome show conserved compartments A and B between 2C and 4C rice, respectively (Figure 3A). We found that rice genome doubling induces switches between compartment A and B (Figure 3A and 3B) with 1% compartments converted from A to B, and 1.39% converted from B to A (Figure 3A). However, we found that the numbers of expressed genes in either conserved compartments (A to A, or B to B) or switched compartments (B to A, or A to B) are of no significant difference between 4C and 2C rice (Figure 3B).

**Figure 3.**
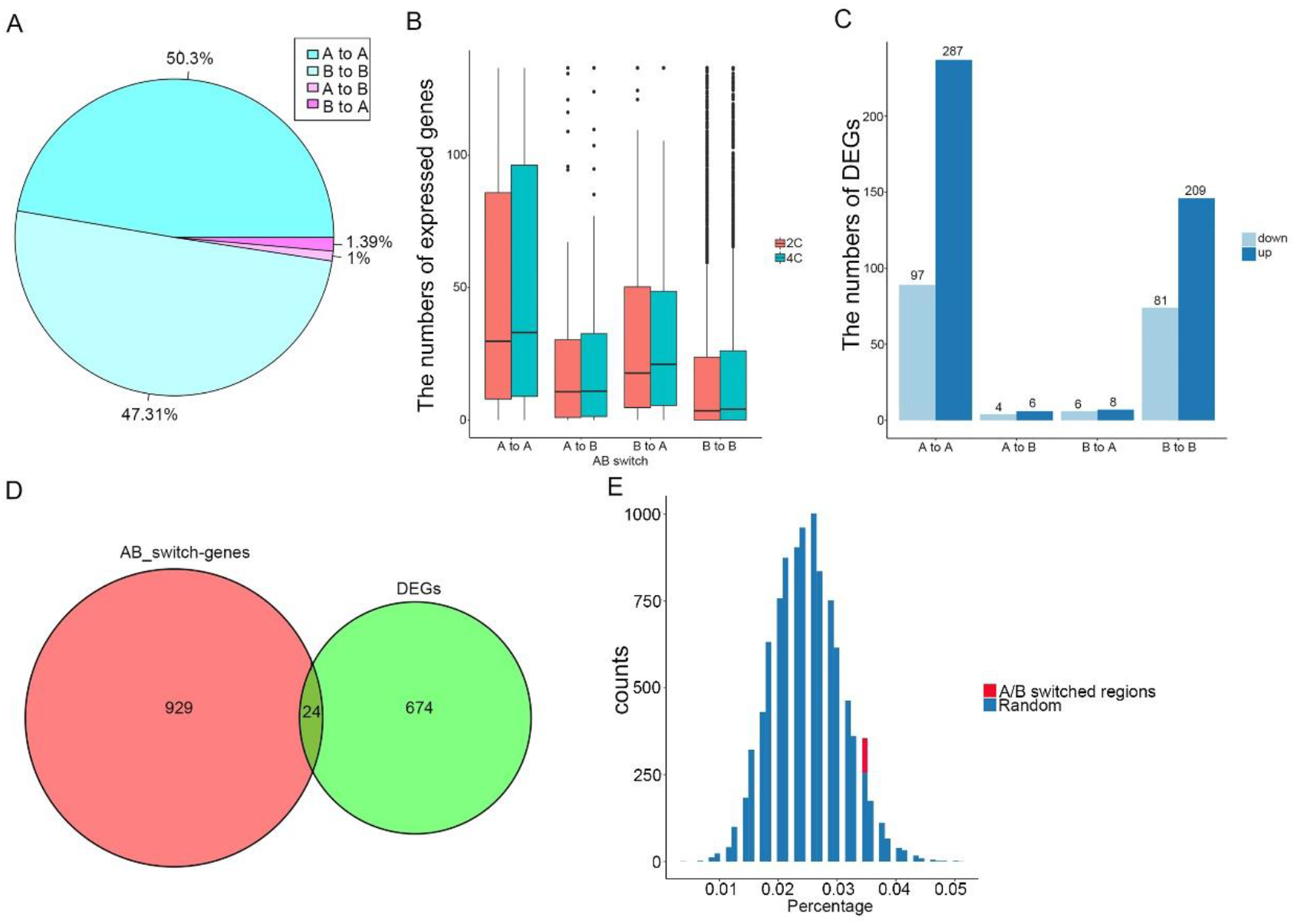
The changed chromatin compartments are uncorrelated with the gene expression. **A**, Pie chart representing the percentages of chromatin compartment switches between 2C and 4C rice. **B**, Boxplots showing the numbers of protein-coding genes in A/B compartments between 2C and 4C rice. **C**, Bar graph showing the statistics of DEGs in the switches of compartments A/B. **D**, Venn diagram showing the numbers of genes in compartment A/B switched regions (pink) and DEGs (green) between 2C and 4C rice. **E**, Histogram of randomly selected DEGs in AB switch regions (n=10000). The red bar chart shows the true AB switch genes in DEGs, and the blue chart shows randomly selected genes with the same number of DEGs overlapped with switched AB compartments. X-coordinate shows proportion of different genes associated with AB switches.

We further analyzed the correlation between chromatin compartment switches and transcription. We found that only 24 DEGs (24/698, about 3.4%) overlapped with switched regions between compartments A and B (Figure 3C and 3D). To know the confidence of the result, we performed bootstrapping randomized analysis. We randomly selected 10000 group of equal number (698) of genes to determine the percentage of those genes overlapped with A/B switched regions. The result showed that 7.24% of the randomly selected groups had more genes than the DEGs overlapped with the switched A/B compartments (Figure 3E), indicating that the overlap of DEGs with the switched A/B compartments is not statistically significant. It means that chromatin compartment switches after rice genome duplication is not related to transcription.

### The differential TAD boundaries and loops are not important for the transcriptional regulation upon rice genome doubling

To know if TAD changes (TADs to non-TAD regions, or non-TADs to TAD regions, are called TAD changes) after rice genome duplication are related to transcription, we first identified TADs by a modified ‘TADcompare’ algorithm (Cresswell and Dozmorov, 2020). 2688 and 2759 TADs were defined in 2C and 4C rice with a median size of 150 kb, respectively (Supplemental Figure S5A). We then categorized and created 5 group TADs with each group having an equal number of TADs. The 5 group TADs were arranged according to gene content with the gene-poorest bin encompasses less than 8 % of all genes while the gene-richest group carries over 50 % of all genes (Figure 4A and 4B).

**Figure 4.**
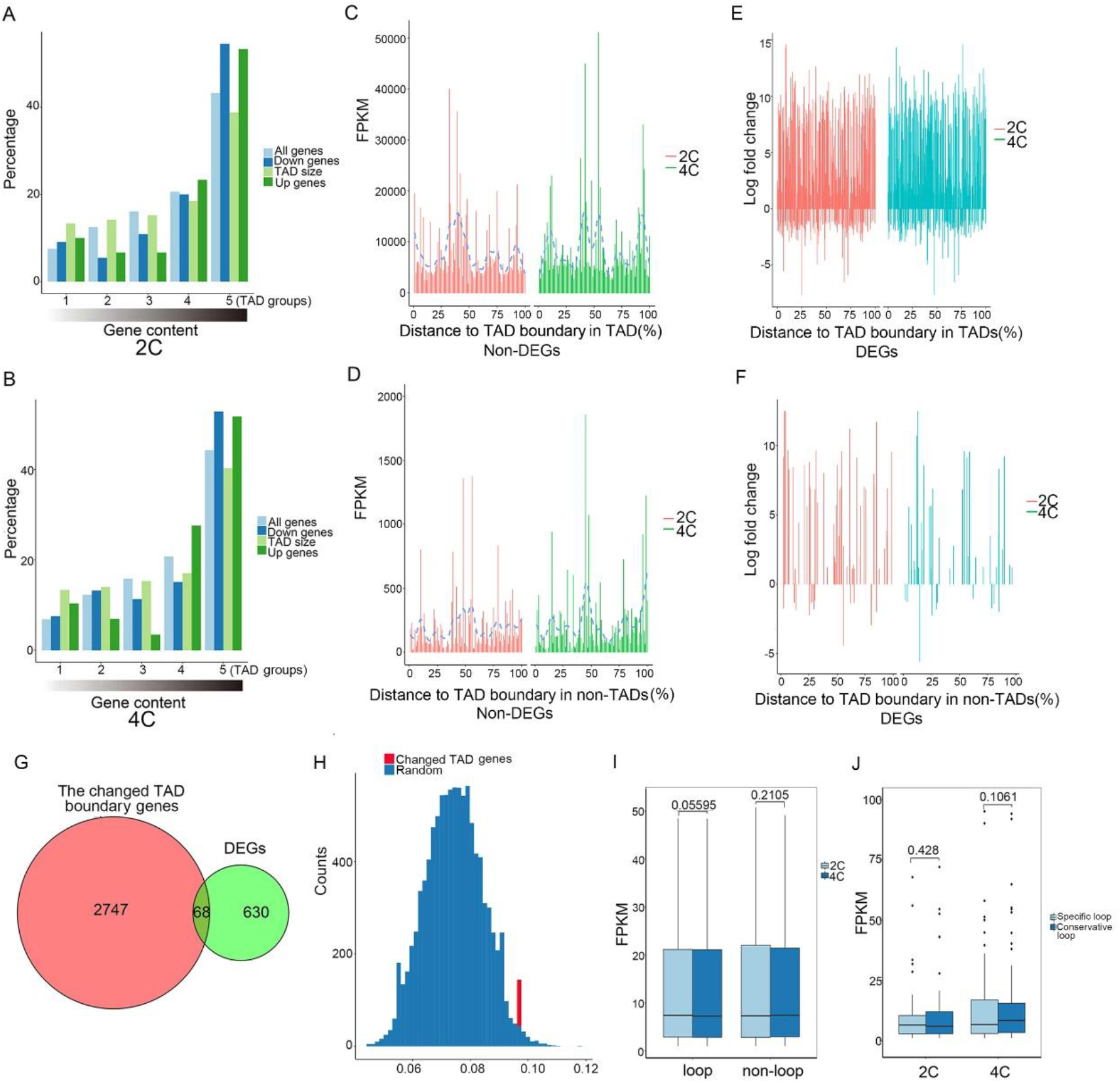
TADs and loops are uncorrelated with gene expression. **A** and **B**, Percentages of genes compared with the percentages of DEGs in 2C rice (A) and 4C rice (B) in TADs grouped by the number of overlapping genes (gene content). The ranking on the x-axis is such that the leftmost group contains 20 % TADs with the lowest number of genes and the rightmost group contains 20 % TADs with the highest number of genes. **C**, Bar plots showing the distance distribution of non-DEGs from the TAD boundary in TADs. X-coordinate shows the percentage from the TSS of genes to the left boundary of TADs, 0 represents the left boundary of TADs, 100 represents the right boundary of TADs, and y-coordinate shows the gene expression. **D**, Bar plots showing the distance distribution of non-DEGs from the TAD boundary in non-TADs. X-coordinate shows the percentage from the TSS of genes to the TAD boundary, and y-coordinate shows the gene expression. **E**, Bar plots showing the distance distribution of DEGs from the TAD boundary in TADs. X-coordinate shows the percentage from the TSS of genes to the TAD boundary, and y-coordinate shows the log fold change of DEGs. **F**, Bar plots showing the distance distribution of DEGs from the TAD boundary in non-TADs. X-coordinate shows the percentage from the TSS of genes to the TAD boundary, and y-coordinate shows the log fold change of DEGs. Non-TAD indicates the regions in the genome except TADs. **G**, Venn diagrams showing numbers of genes in changed TADs (pink) and DEGs (green) between 2C and 4C rice. **H**, Histogram of randomly selected DEGs in TAD changed regions (n=10000). The red bar chart shows the true TAD changed genes in DEGs, and the blue chart shows the TAD changed genes in random number of DEGs. X-coordinate shows proportion of different genes associated with TAD changed. TAD changed means that a gene goes from TAD to non-TAD, or non-TAD to TAD, after chromosome doubling. **I**, Boxplots showing the normalized RNA-seq FPKMs between genes in loops and non-loops in 2C and 4C rice. **J**, Boxplots showing the normalized RNA-seq FPKMs of genes in specific loops and conservative loops in 2C and 4C rice. Specific loops indicate loops unique to 2C or 4C, conservative loops indicate loops shared by 2C and 4C.

In addition to TADs, we defined the 5 kb regions adjacent to TADs as TAD boundaries, and other regions as non-TADs, given that the TAD structures were calculated at a 10 kb resolution. In both 2C and 4C rice, the overall protein-coding gene density in TAD regions of 4C rice was similar to that of 2C rice (Supplemental Figure S5B), and the overall protein-coding gene density in non-TAD regions of 4C rice was slightly higher than that of 2C rice near TAD boundaries (Supplemental Figure S5C). The relative locations of non-DEGs altered in 2C rice compared to 4C rice in both TADs and non-TADs (Figure 4C and 4D). In contrast, the relative locations of DEGs in both TADs and non-TADs in 4C rice are similar to those in 2C rice (Figure 4E and 4F), indicating that the changes of TADs during rice genome duplication affect the relative locations of some non-DEGs, but did not cause significant differential expression. Among 698 DEGs in total, 628 locate in TADs of 2C rice and 652 in TADs of 4C rice (Supplemental Figure S6). Only a small portion of DEGs (68/698, about 9.76%) localizes in TAD changed regions between 2C and 4C rice genome (Figure 4G and Supplemental Data Set S4). We then performed bootstrapping randomized analysis. We randomly selected 10000 group of equal number (698) of DEGs to determine the percentage of those genes overlapped with TAD changed genes, the result showed that the percentage of DEGs (9%) localized in the percentage of randomly selected control genes (~12%) (Figure 4H), indicating that the changed TAD boundaries after rice genome duplication is not related to transcription.

To know whether loop changes after rice genome duplication are related to transcriptional regulation, we first annotated 4822 loops in 2C rice and 5365 loops in 4C rice and identified 79 loops specific for 2C rice and 81 loops specific for 4C rice (Supplemental Figure S5D and 5E; Supplemental Data Set S5). We found that FPKMs of genes in loops or non-loops between 2C and 4C rice are of no significant difference (Figure 4I), and that FPKMs of genes in specific loops and conservative loops are also of no significant difference in both 2C and 4C rice (Figure 4J), suggesting that the chromatin loops are not important for the transcriptional regulation when 2C rice is duplicated to 4C rice.

### DNA methylations in the promoter regions of genes in compartment A/B switched regions and TAD boundaries are not involved in transcriptional regulation upon rice genome duplication

It was known that the global level of DNA methylation increases when 2C rice is duplicated to 4C rice (Zhang et al., 2015). We analyzed DNA methylation in different regions of genes and the relationship between DNA methylation and 3D genomic structures in 2C and 4C rice. The genome-wide DNA methylation revealed by WGBS-seq (Supplemental Data Set S6) indicate that the average levels of DNA methylations in CG, CHG and CHH contexts are increased in 4C rice compared to 2C rice in genes, upstream and downstream of genes with each gene defined by a continuous exon and intron sequence (Supplemental Figure S6A). In 2C and 4C rice, 59% and 62.5% of DNA methylation are in CG context, 29% and 31% in CHG context, and 3.5% and 3.5% in CHH context, respectively (Supplemental Figure S6B). Compared to 2C rice, 4C rice exhibits increased proportions of methylated cytosines in CG and CHG contexts (Supplemental Figure S6B), and CG methylation level is higher than CHG and CHH methylations (Supplemental Figure S6B), similar to the previous report (Zhang et al., 2015).

As DNA methylation in the regulatory regions of genes plays an important role in transcriptional regulation (Zhang et al., 2018b), we compared the DNA methylation in DEG promoters between 2C and 4C rice. All DEGs and non-DEGs were classified according to log fold change (LFC) of genes (Supplemental Figure S7). The results showed that there was no significant difference of DNA methylation levels in DEG promoters between 2C and 4C rice, and the CG and CHG methylation levels of non-DEG promoters in 2C rice are lower than those in 4C rice (Supplemental Figure S7).

We then analyzed the correlation between DNA methylation in 4 kb upstream of genes in compartment A/B switched regions and the transcriptional changes of these genes upon rice genome duplication. We found that the CG, CHG and CHH methylation levels in 4 kb upstream of all genes in compartment A/B switched regions are not significantly changed in 4C rice compared to those in 2C rice (Figure 5A). Moreover, DNA methylation levels in 4 kb upstream of DEGs in compartment A/B switched regions are similar to those of non-DEGs in both 2C and 4C rice (Figure 5B and 5C). These results suggested that there is not an obvious link between DNA methylation in the upstream of genes in compartment A/B switched regions and the transcriptional regulation of these genes upon rice genome doubling.

**Figure 5.**
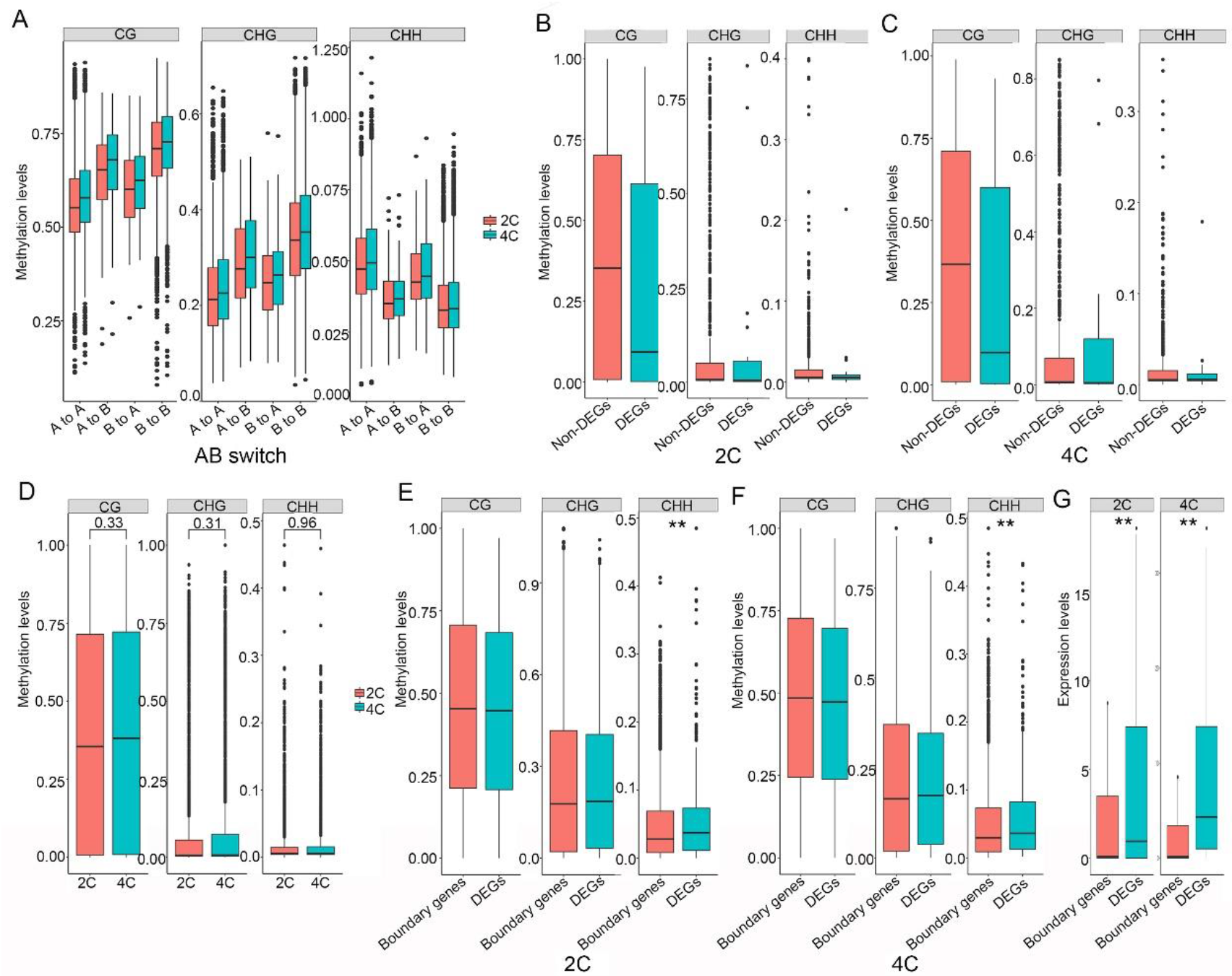
The DNA methylation levels show no changes between DEGs and non-DEGs or between 2C and 4C rice in the compartment A/B switched regions. **A**, Comparisons of CG, CHG and CHH methylation levels in 4 kb upstream of genes in the compartment A/B switches between 2C and 4C rice. **B**, Comparisons of CG, CHG and CHH methylation levels in 4 kb upstream of DEGs with non-DEGs in the compartment A/B switches of 2C rice. **C**, Comparison of methylation levels of CG, CHG and CHH contexts in 4 kb upstream of DEGs with non-DEGs in the compartment A/B switches of 4C rice. **D**, Comparison of CG, CHG and CHH methylation levels in 4 kb upstream of TAD boundary genes (3247) between 2C and 4C rice. **E**, Comparison of CG, CHG and CHH methylation levels between TAD boundary genes and DEGs in 2C rice. **F**, Comparison of CG, CHG and CHH methylation levels between 4 kb upstream of TAD boundary genes and DEGs in 4C rice. \**p* value < 0.05; \*\**p* value < 0.01. **G**, Boxplots showing FPKMs between TAD boundary genes and DEGs in 2C and 4C rice. The *p* values were tested by Wilcoxon–Mann–Whitney test.

To know the role of the methylation of TAD boundary genes in the transcriptional regulation, we compared the methylation level of TAD boundary genes with that of DEGs upon rice genome doubling. We found that the CG, CHG and CHH methylation levels in 4 kb upstream of TAD boundary genes (3247) in 4C rice remain unchanged compared to those (3203) in 2C rice (Figure 5D; Supplemental Data Set S7). In addition, CG and CHG methylation levels in 4 kb upstream of TAD boundary genes do not change significantly compared to those in 4 kb upstream of DEGs in both 2C and 4C rice (Figure 5D). Moreover, the CHH methylation level in 4 kb upstream of TAD boundary genes is lower than that of DEGs in both 2C and 4C rice (Figure 5E and 5F). In contrast, FPKMs of TAD boundary genes are lower than those of DEGs (Figure 5G), which is opposite to the expectation that FPKMs of the hypomethylated TAD boundary genes are higher than FPKMs of the hypermethylated DEGs. We concluded that DNA methylation in the upstream regions of TAD boundary genes is not involved in transcriptional regulation.

We further analyzed the relationships between DEGs and differential methylation regions (DMRs) which were known to participate in gene transcription (Schmitz et al., 2011). Totally, 1484 CG, 89 CHG, and 1 CHH DMRs were identified between 2C and 4C rice, including 892 CG, 26 CHG, and 1 CHH hypermethylated DMRs, and 592 CG, 63 CHG, and 0 CHH hypomethylated DMRs. We then examined the genomic distances between DEGs and DMRs (Becker et al., 2011; Schmitz et al., 2011). The distance between the TSS locus of each DEG and all DMRs was calculated, and the shortest distance was taken as the distance between a given DEG and DMRs. We found that the genomic distances between DEGs and DMRs are very long (Supplemental Figure S8), suggesting that there are no obvious relationships between DEGs and DMRs.

### Hypermethylated TEs across non-DEGs in compartment A/B switched regions correlate with the inhibited gene transcription upon rice genome doubling

Given that the number, distance and methylation level of TEs were known to affect the expression of genes neighboring the TEs in *Arabidopsis*, rice and maize (Wang et al., 2013; Zhang et al., 2015; Forestan et al., 2017), we addressed if DNA methylation in TEs is involved in 3D genome-mediated transcriptional regulation. In rice 9311 genome, 14.57% of TEs localize in genes, 38.19% in upstream of genes, and 38.33% in downstream of genes. We compared the methylation levels in CG, CHG, and CHH contexts between 2C and 4C rice for 12 major types of TEs including class I retrotransposons Copia, Gypsy, LTR, LINE and SINE, and class II transposons Helitron, Stowaway, DNA, Harbinger, MULE_MuDR and hAT. The results indicate that the methylation levels of Copia, DNA, Harbinger, LINE, MULE_MuDR, and SINE in 4C rice are different from those in 2C rice (Supplemental Figure S9 and 10). We then compared the methylation levels of TEs in gene promoters between 2C and 4C rice. The results showed that there was no significant difference of methylation levels in TEs in promoters of DEGs and non-DEGs between 2C and 4C rice (Supplemental Figure S11).

To analyze the effect of DNA methylation in TEs across genes in compartment A/B switched regions on gene expression, we compared the methylation levels of all TEs, Class I and II TEs across non-DEGs with those across DEGs in compartment A/B switched regions between 2C and 4C rice, with TEs across a gene defined by TEs in the gene body and 4 kb regions flanking the gene. The results indicate that all TEs and Class I TEs in non-DEG bodies show hypermethylation in CG, CHG and CHH contexts (Figure 6A and 6B), and Class II TEs in non-DEG bodies show no change in CG, CHG and CHH contexts compared to those in DEG bodies in 2C and 4C rice (Figure 6C). In addition, the results showed that all TEs in 4 kb regions flanking non-DEGs show hypermethylation in CG context (Figure 6D), Class I TEs show hypermethylation in CG and CHG contexts (Figure 6E), and Class II TEs show no change of methylation in CG, CHG and CHH contexts compared to those in 4 kb regions flanking DEGs in 2C and 4C rice (Figure 6C). In contrast, FPKMs of non-DEGs in compartment A/B switched regions are lower than those of DEGs in both 2C and 4C rice (Supplemental Figure S12). These results show that the hypermethylation of TEs across non-DEGs in compartment A/B switched regions correlates with the suppressed transcription of these genes.

**Figure 6.**
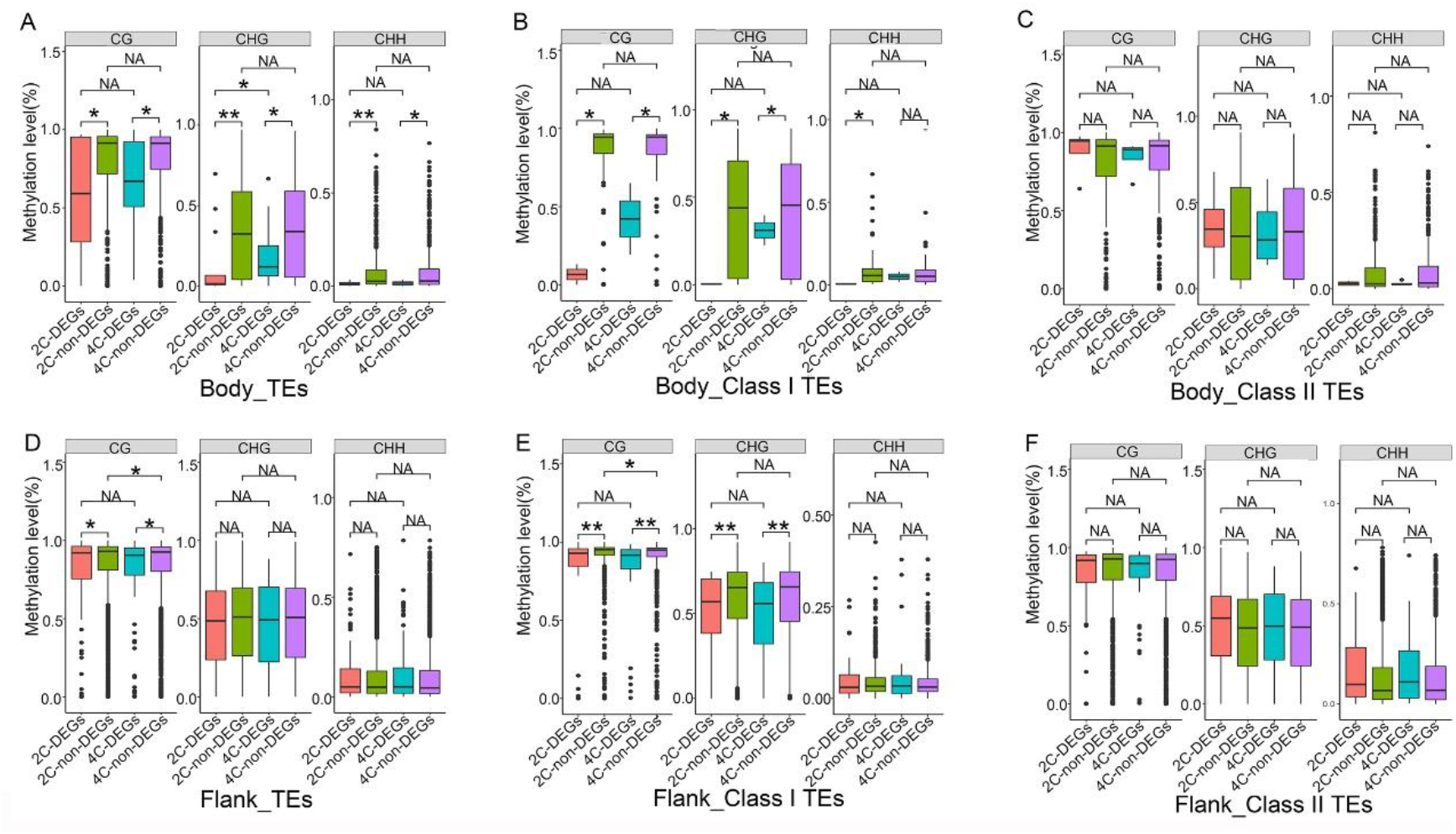
TEs in bodies of non-DEGs or regions flanking non-DEGs in compartment A/B switched regions are hypermethylated compared to those of DEGs in 2C and 4C rice. **A**, Comparison of TE methylation levels in gene bodies between non-DEGs and DEGs in compartments A/B switched regions of 2C and 4C rice. **B**, Comparison of Class I TE methylation levels in gene bodies between non-DEGs and DEGs in compartments A/B switched regions of 2C and 4C rice. **C**, Comparison of Class II TE methylation levels in gene bodies between non-DEGs and DEGs in compartments A/B switched regions of 2C and 4C rice. **D**, Comparison of TE methylation levels in regions flanking genes between non-DEGs and DEGs in compartments A/B switched regions of 2C and 4C rice. **E**, Comparison of Class I TE methylation levels in regions flanking genes between non-DEGs and DEGs in compartments A/B switched regions of 2C and 4C rice. **F**, Comparison of Class II TE methylation levels in regions flanking genes between non-DEGs and DEGs in compartments A/B switched regions of 2C and 4C rice. (Class I is retrotransposons including Copia, Gypsy, LTR, LINE and SINE, and class II is transposons including Helitron, Stowaway, DNA, Harbinger, MULE_MuDR and hAT).

### Hypermethylated CG in TEs across TAD boundary genes correlates with the suppressed gene transcription upon rice genome duplication

As TAD boundary plays an important role in the regulation of local transcription and epigenetic landscape in different species (Bonev et al., 2017; Yang et al., 2019; Ouyang et al., 2020), we evaluated the impacts of the methylation levels of TEs across TAD boundary genes on the role of 3D chromatin structure in transcriptional regulation. As TEs including Copia, DNA, Harbinger, LINE, MULE_MuDR and SINE exhibit differential DNA methylation between 2C and 4C rice (Supplemental Figure S9 and 10), we compared the methylation levels of these TEs across TAD boundary genes in 2C and 4C rice to those across DEGs, including the gene bodies and 4 kb regions flanking these genes. The results indicate that Copia, DNA and MULE-MuDR across TAD boundary genes show hypermethylation in CG context, no change in CHG context, and hypomethylation in CHH context compared to those across DEGs in 2C and 4C rice (Figure 7A–7D).

**Figure 7.**
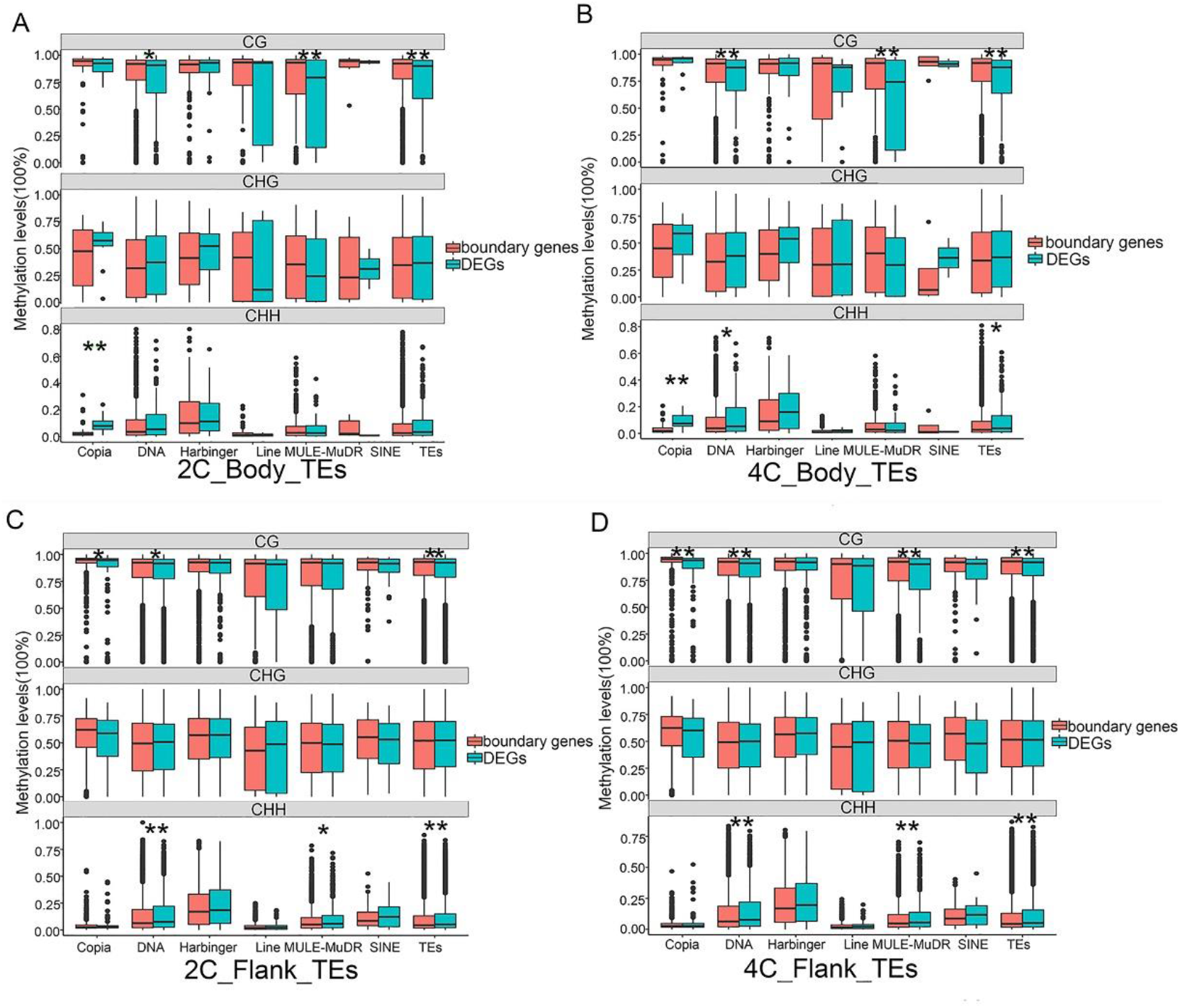
TEs across non-DEGs in TAD boundaries are hypermethylated compared to those across DEGs in 2C and 4C rice. **A**, Comparison of TE methylation levels in gene bodies between TAD boundary genes and DEGs in 2C rice. **B**, Comparison of TE methylation levels in gene bodies between TAD boundary genes and DEGs in 4C rice. **C**, Comparison of TE methylation levels in regions flanking genes between TAD boundary genes and DEGs in 2C rice. **D**, Comparison of TE methylation levels in regions flanking genes between TAD boundary genes and DEGs in 4C rice. (“flanking” represents the 4 kb regions flanking genes).

To verify the CHH hypomethylation in TEs, we analyzed the clusters of siRNAs, as siRNAs often mediate CHG and CHH types of methylation in plants, and 24 nt siRNAs can enter the RdDM pathway to trigger DNA methylation and transcriptional silencing to suppress TE activities (Matzke and Mosher, 2014). We investigated the relationship between CHH methylation level and siRNA abundance in TEs across TAD boundary genes and DEGs. We gained sRNA length profiles similar to previously reported (Peng et al., 2011; Zhang et al., 2015) in 2C and 4C rice (Supplemental Figure S13A). In total, 154,968 siRNA clusters were identified in 2C rice, and 147,589 clusters in 4C rice. Most siRNA clusters overlapped with class I and class II TEs, and the cluster fractions in Class I TEs, Class II TEs, genes and intergenic regions of 4C rice are similar to those of 2C (Supplemental Figure S13B). The lower CHH methylation level in TEs across TAD boundary genes than that in TEs across DEGs (Figure 7A–7D) is parallel with the lower siRNA level in TEs across TAD boundary genes than that in TEs across DEGs (Supplemental Figure S14A-14D).

The CG methylation level of TEs across TAD boundary genes is much higher than those of CHG and CHH methylations in 2C and 4C (Supplemental Figure S15A and 15 B; Figure 7A–7D). In contrast, FPKMs of TAD boundary genes are lower than FPKMs of DEGs in 2C and 4C rice (Figure 5G), indicating that CG hypermethylation of TEs across TAD boundary genes correlates with the inhibited transcriptions of these genes.

### Hypermethylated TEs adjacent to TAD boundary genes suppress the transcription of TAD boundary genes upon rice genome doubling

Next, we confirmed the role of TE methylation in the regulation of TAD boundary genes by comparing the expression levels between TAD boundary genes and DEGs at different distances to the closest TE in 2C and 4C rice. As more than 97% TEs are clustered in the 1.2 kb regions away from TAD boundary genes or DEGs in both 2C and 4C, we analyzed the expressions of TAD boundary genes or DEGs from 0 to 1.2 kb away from TEs (Supplemental Data Set S8). In addition, the average TE length is 337bp, therefore, 400bp was chosen as the distance interval. Compare to TEs in body of boundary genes (0 bp in Figure 8A for 2C; Figure 8B for 4C), CG, CHG and CHH methylation levels in TEs at 0-400 bp increase slightly (0-400 bp in Figure 8A for 2C; Figure 8B for 4C), while the expression level of TAD boundary genes deceases slightly (0-400 bp in Figure 8C for 2C; Figure 8D for 4C); Compare to TEs at 0-400 bp, CG, CHG and CHH methylation levels of TEs at 400-800 bp decrease (400-800 bp in Figure 8A for 2C; Figure 8B for 4C), while the expression level of TAD boundary genes increases dramatically (400-800 bp in Figure 8C for 2C; Figure 8D for 4C); Compare to TEs at 400-800 bp, CG, CHG and CHH methylation levels of TEs at 800-1200 bp increase (400-800 bp in Figure 8A for 2C; Figure 8B for 4C), while the expression level of TAD boundary genes decease (400-800 bp in Figure 8C for 2C; Figure 8D for 4C). Importantly, the trends of methylations in TEs adjacent to TAD boundary genes are opposite to the trends of gene expressions in both 2C and 4C rice (Figure 8A *vs*. 8C and Figure 8B *vs*. 8D). The troughs of DNA methylation levels appear within 400-800bp away from the closest TE (Figure 8A and 8B). These data suggested that the expression levels of TAD boundary genes are positively correlated with the distance to the closest TE in 2C and 4C rice.

**Figure 8.**
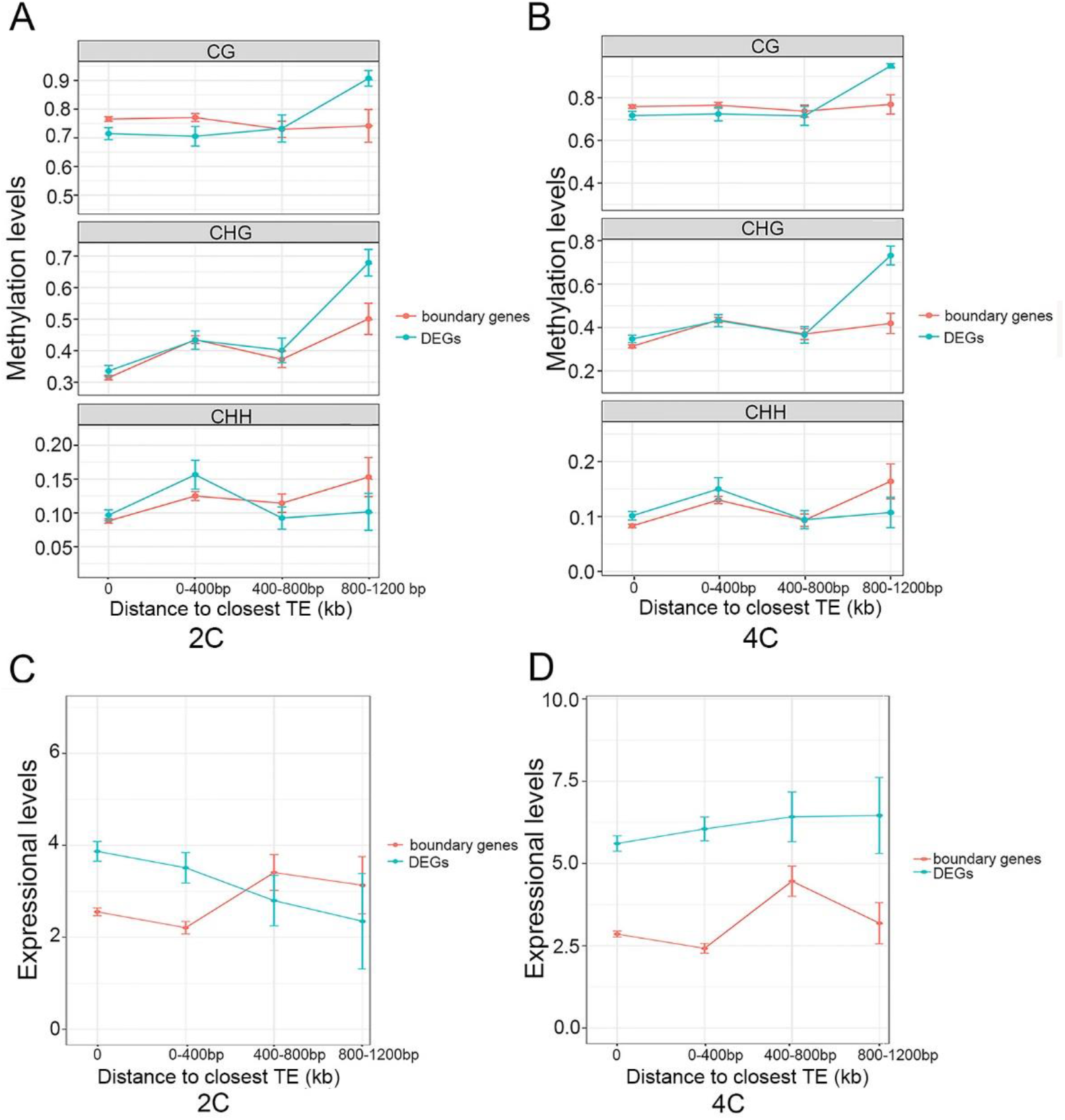
The distances from TEs to genes are related to the expression levels of these genes. **A**, Comparison of TE methylation levels related to the distance to the closest TE between TAD boundary genes and DEGs in 2C rice. **B**, Comparison of TE methylation levels related to the distance to the closest TE between TAD boundary genes and DEGs in 4C rice. “0” indicates genes overlapped with TEs in body regions. Error bars indicate SEMs. **C**, Comparison of gene expression levels related to the distance to the closest TE between TAD boundary genes and DEGs in 2C rice. **D**, Comparison of gene expression levels related to the distance to the closest TE between TAD boundary genes and DEGs in 4C rice.

In contrast, the expression levels of DEGs of 2C rice (2C *vs*. 4C) decrease with an increasing distance from TEs. Conversely, the expression levels of DEGs of 4C rice (4C *vs*. 2C) increase with an increasing distance from TEs, and the peak of those expressional levels appear within 800-1200bp away from the closest TE (Figure 8C and 8D). Importantly, the trends of methylation in TEs adjacent to DEGs within 1.2 kb are not opposite to that of gene expression in both 2C and 4C rice (Figure 8A *vs*. 8C and Figure 8B *vs*. 8D), which indicate that the expression levels of DEGs are uncorrelated with the distance to the closest TE within 1.2 kb in 2C and 4C rice. We proposed that the hypermethylation in TEs adjacent to TAD boundary genes may buffer the impact of TAD boundaries on gene transcription.

## Discussion

Polyploidization promotes the evolution of higher plants (Wendel, 2000; Otto, 2007). Many plants, including *Arabidopsis*, rice, soybean, poplar, sorghum, and maize, might have experienced whole-genome duplication events during their evolution (Jiang et al., 2013). More attention has been paid to the phenotypic, epigenetic and gene expression changes of autopolyploidy plants, while less attention to the changes of 3D genome and their impacts on gene expression during autopolyploidization. The study of 3D chromatin topology of autopolyploid crops is important for understanding the contribution of spatial organization of genome to the success of polyploidy species. We previously reported that the altered chromatin interactions in 4C dicotyledonous *Arabidopsis*, compared to its 2C progenitor, modulate the transcription (Zhang et al., 2019). The study on soybean also proved that chromatin loop reorganization was involved in gene expression divergence during soybean domestication (Kato et al., 2020). To better understand the change of 3D genome during monocotyledonous genome doubling and its potential effect on gene expression, we performed Hi-C, epigenome and transcriptome analysis in 4C rice and its progenitor 2C rice. Our results showed that rice genome doubling obviously dampens intra-chromosomal interactions. Importantly, the changes of 3D chromatin structure upon rice genome duplication were found to be uncoupled with gene transcriptions, which is reminiscent of several reports that 3D chromatin structure are unrelated to gene expression (Espinola et al., 2021; Ing-Simmons et al., 2021), (Dong et al., 2020).

Although DNA methylation variation has been observed (Lee and Chen, 2001; Madlung et al., 2002; Wang et al., 2004; Lukens et al., 2006; Kenan-Eichler et al., 2011) during genome duplication in plants, moreover, TE methylations were also known to be able to affect the expression of nearby genes in *Arabidopsis* (Wang et al., 2013), rice (Zhang et al., 2015), and maize (Forestan et al., 2017), however the specific role of TE methylation in 3D chromatin structure alteration is not clear. It has been known that the 3D genome architecture modulated gene transcription by bringing together distant promoter, enhancer, and other cis-regulatory elements (Spitz and Furlong, 2012). Rice genome doubling might be accompanied by the locational changes of regulatory elements such as promoters and enhancers, which, however, do not result in the change of gene expression. Interestingly, we found that the disconnection between transcriptional regulation and A/B switches or TAD boundaries upon genome duplication is not due to alterations of DNA methylation in the regulatory sequences of the A/B switch- and TAD boundary-related genes, but to the changes of DNA methylation in TEs adjacent to these genes. Our study suggested that autopolyploidization may stimulate TE modification to reduce the effect of the changes of 3D chromatin structure on gene expressions during genome doubling, the underlying mechanism might be that the decrease of intra-chromosomal interactions is beneficial to the activities or methylation of TEs by increasing the genomic accessibility to DNA methyltransferases and/or demethylases to antagonize the effect of decreased chromatin interactions on genomic regulation, resulting in disruption of the correlation between 3D chromatin structure and gene expression, which might contribute to the success of polyploidy plants during evolution.

In human and animals, many CTCF binding sites are derived from TEs (Schmidt et al., 2012; Trizzino et al., 2017) and CTCF protein defines the TAD boundaries to mediate the formation of TADs (Dixon et al., 2012). We found that TEs adjacent to TAD boundary genes can inhibit the expression of these genes in both 4C and 2C rice. However, CTCF-like proteins have not been identified in plants, possibly, there are other elements or proteins in plants that might play a similar role in defining TAD boundaries to that of CTCF factor in animals.

Increased inter-chromosome interactions and decreased intra-chromosome arm interactions were observed in 4C *Arabidopsis* compared to its 2C progenitor (Zhang et al., 2019). 4C *Arabidopsis* seedlings show obvious phenotypic changes in vegetative stage, including serrated leaves, more rosette leaves, and increased whole plant size (Zhang et al., 2019), which is related to the alteration of 3D chromatin structure, resulting in changes the gene expression at this stage. In contrast, no obvious phenotypes of monocotyledons 4C rice were observed at vegetative stage, which might be related to that the 3D chromatin structural changes are not related to gene expression when 2C rice is duplicated to 4C rice. In ripening stage, the 4C rice acquires many morphologic traits compared to 2C rice (Zhang et al., 2015) (Figure 1), it will be therefore of interest to study the relationship between transcriptional regulation and 3D chromatin organization in the reproductive stage upon rice genome duplication. In addition, there are 698 DEGs in vegetative stage, since their differential expressions are not caused by DNA methylation alteration or 3D chromatin rearrangement, there may be other epigenetic mechanisms which are involved in modulating the expressions of these 698 genes, such as histone modification, acetylation or noncoding RNAs.

The relationships between higher-order chromatin structure with other epigenetic regulation, including DNA methylation, histone modifications and noncoding RNAs are implicated at multiple developmental processes. In human cells, it was found that large DNA methylation nadirs can mediate the formation of long loops (Zhang et al., 2020). In *Arabidopsis*, it was reported that lncRNA (APOLO) can modulate local chromatin 3D conformation through regulating the conformation of DNA-RNA loops within the nuclei (Ariel et al., 2020). Histone modifications mediate the impact of genetic risk variants related to schizophrenia by modulating chromatin higher-order structure (Punzi et al., 2018). Here we found that TE methylation diminishes the impact of the alteration of chromatin higher-order structure on gene expression. It will be worthy to study the specific epigenetic networks that include high-order chromatin architecture, DNA and histone modifications, noncoding RNAs and other epigenetic factors during plant genome duplication.

## Materials and Methods

### Plant materials

Autotetraploid (4C) rice line was artificially synthesized from *O. sativa ssp. indica* cultivar 9311 (2C).

### Characterization of agronomic traits

All plants were cultured in nutrient solution (Han LZ, 2006) in a growth chamber with 28°C /25°C (day/night) and 12 h/12 h (light/dark) cycles. After germination for 15 days, the rice seedlings were transferred to the field. The agronomic traits including plant height, flag leaf length, flag leaf width, tillering number, panicle length, grains per panicle, grain weight, grain length, grain width were scored in parallel between 2C and 4C rice. The traits were selected and analyzed according to the Descriptors and Data Standard for Rice (*O. sativa* L.) (Han LZ, 2006).

### Flow cytometry

Ten-day-old 2C and 4C rice seedlings were collected into pre-cooled plate, and then chopped with a new razor blade to release nuclei in the sterile lysis buffer (45 mM MgCl_2_·6H_2_O, 30 mM trisodium citrate, 20 mM MOPS, 1% Triton X-100, pH 7.0) for 3-5 min until the buffer turns green. Transfer the mixture into strainer. The filtrates were added with the final concentration of 1 ng / ml DAPI solution, and then cultured in dark on ice for 30 min. The ploidy levels were measured by flow cytometry (Beckman Coulter MoFlo XDP, USA).

### Hi-C library preparation

Hi-C experiments were performed essentially as described (Grob et al., 2014) with some modifications. Two biological replicates of 2C and 4C rice were performed. In short, 2.5g of the above ground parts of 10-day-old seedlings were fixed (2% formaldehyde, 10% PBS) and ground into powder in liquid nitrogen. The extracted nuclei were digested by incubation with 600 U *Hin*dIII restriction enzyme at 37°C overnight, and the digested chromatin at 1μl 10 mM dATP, dTTP, dGTP and 25μl 0.4mM biotin-14-dCTP and 100 U Klenow fragment was placed at 37°C for 45 min. The ligation reaction was then carried out in 10× volume of ligation buffer and shaken with 745 μl 10× ligation buffer, 10% Triton X-100, 80μl 10 mg/ml BSA and ATP, 100 Weiss U T4 DNA ligase at 16°C for 6 hours. Then reversely cross-linked with proteinase K at 65°C overnight. Subsequently, the extracted chromatin was fragmented into an average size of 300 bp by ultrasound (Covaris s220). The Hi-C library was constructed with NEB Next Multiplex Oligos kit and KAPA Hyper Prep Kit. The final library was sequenced on Illumina HiSeq X Ten instrument with 2 × 150 bp reads.

### Hi-C sequencing data processing

Hic-pro (Servant et al., 2015) and Bowtie2 (Langmead and Salzberg, 2012) were used for Hi-C read mapping. The clean Hi-C reads of 2C and 4C rice were aligned to the genome of *O*. *sativa* Indica (Yu et al., 2002) after removing the adapter. Following processing with HiC-Pro and Juicer software (Durand et al., 2016), valid pairs of 2C and 4C rice were used to create interaction matrixes with bin size 50 kb for further analysis. The reproducibility of two biological replicates was tested with Pearson correlation coefficient from the ICE normalized interaction matrixes (Lin et al., 2018). Hicpro2juicebox was used to generate input file for Juicebox. The interaction matrixes were normalized with KR method from Juicer at resolutions 5 kb, 10 kb and 50 kb (Durand et al., 2016). After excluding the pericentromeres as reported (Grob et al., 2014), the first principal component was used to identify compartments with Juicer at 50 kb resolution, and the direction of compartment with high gene expression was defined as A compartment, and the opposite direction as B compartment. We calculated loops and differential loops at 5 kb and 10 kb resolutions, using HICCUPS and HICCUPS Diff in Juicer (Durand et al., 2016). We used TADCompare (Cresswell and Dozmorov, 2020) to calculate the TAD structure at a 10 kb resolution.

### Calculation of chromatin interactions and interaction decay exponents

The normalized interaction matrix of 2C rice was divided by the normalized interaction matrix of 4C rice, and all zeros in the matrix were replaced with the smallest non-zero elements in each matrix to analyze the difference between 2C and 4C rice interaction matrices. We used log2 transform and median normalization to standardize the difference matrix. Interaction decay exponents (IDEs) of chromosomes, pericentromeres and telomeres were calculated (Grob et al., 2014) to study the variation of interaction frequency dependent on the genome distance, the CPM normalization method was used to process the data.

### Bootstrapping analysis

In the bootstrapping strategies (Buonaccorsi et al., 2018), we randomly selected 10000 groups (n = 10000 times) of the same number of DEGs, and performed the same analysis to determine the percentage of those genes overlapped with A/B switch region genes or TAD changed genes. When the percentile of the test sample was higher than the top five percentiles of the control distribution, it is considered as statistically significant.

### RNA-seq analysis

Total RNAs were extracted from 10-day-old 2C and 4C rice seedlings using the RNeasy plant mini kit (Qiagen). cDNA library construction and sequencing were carried out by Beijing Genomics Institute (BGI) using BGISEQ-500 platform for 50 bp single-end sequencing as previously described (Huang et al., 2018). At least 20 M clean reads of sequencing depth were obtained for each sample. Three independent biological replicates were performed. The clean reads were separately aligned to the genome of *O. sativa* Indica (Yu et al., 2002) with orientation mode using Tophat software (http://tophat.cbcb.umd.edu/). The fragments per kilobase of exon per million mapped reads (FPKMs) method was used to calculate the expression level of each transcript. The differential expression analysis was carried out using the classical normalization method of DESeq2 R package (Love et al., 2014) with a 0.05 *p*-value, 0.05 false discovery rate, and cutoff of 1 log-fold change. The hypergeometric test was performed as previously described (Wollmann et al., 2017). Blast2GO method was used to find homologous genes in japonica rice genome (MSU), and GO functional enrichment analysis was performed by DAVID (https://david.ncifcrf.gov/).

### DNA methylation analysis

For whole genome bisulfite sequencing (WGBS), genomic DNA (gDNA) was extracted from 10-day-old 2C and 4C rice seedlings with the DNeasy plant mini kit (Qiagen) per manufacturer’s introduction. Library construction and sequencing were performed by Beijing Genomics Institute (BGI) using Illumina HiSeq-2000 for 100 bp paired-end sequencing. To facilitate the analysis of DNA methylation data, we used Batmeth2 (https://github.com/GuoliangLi-HZAU/BatMeth2), an integrated multi-functional software for DNA methylation analysis (Zhou et al., 2019a), including sequencing sequence quality filtering, DNA methylation sequence alignment, DNA methylation level calculation and functional annotation.

### Calculation of the distance between a DEG and differentially methylated regions (DMRs)

The whole genome was divided into 1000-bp bins to identify the DMRs in which the absolute value of difference in DNA methylation between 2C and 4C was 0.6 or above, and the adjusted *q* value of Fisher’s exact test was 0.05 or less by using BatMeth2 (Zhou et al., 2019b). Finally, the shortest genomic distance of a given DEG and all DMRs in turn was calculated.

### TE annotation and analysis

By running RepeatMasker (v4.0.3, www.repeatmasker.org), the repetitive library of RepBase (v20130422) was used to compare the rice reference genome sequences. To compare the methylation status of TEs between the 2C and 4C rice genomes, we excluded TEs with less than 40% of cytosines and coverage of BS reads less than 3. The remaining TEs were used for further analysis. Using this cut-off value, we obtained a data set of 478599 TEs for the subsequent analysis.

### Small RNA-seq and data processing

Small RNAs (sRNAs) were isolated from 10-day-old rice seedlings using mirVana^™^miRNA Isolation Kit (Ambion, AM1561) and sequenced by Illumina high-throughput sequencing. The small RNA data were processed and analyzed according to the previous description (McCormick et al., 2011) with minor modifications. In brief, the raw sequencing reads were trimmed using cutadapt (v1.2.1) to remove adapters, and sRNAs between 16 and 35 nt in lengths were selected and mapped to the rice genome (Yu et al., 2002; Zhao et al., 2004).

sRNAs that matched against the databases including the Rfam database (Burge et al., 2013) and miRBase (Kozomara and Griffiths-Jones, 2011) were discarded. 24-nt reads that did not match miRNAs, snRNAs, rRNAs, tRNAs, or snoRNAs were filtered and mapped to the genome 1–1,000 times as siRNAs for analyses. The siRNA count was based on the total abundance of genome matched small RNA reads, normalized to reads per million, excluding sRNAs of the above structures, and dividing the number of reads evenly by the number of genome hits. A siRNA cluster was defined as containing at least five different siRNA reading sequences, and adjacent reading sequences less than 200 bp apart were combined into a cluster.

### Accession numbers

The Hi-C, WGBS and RNA-seq datasets have been submitted to NCBI (PRJNA725914).

## Supplemental data

The following materials are available in the online version of this article.

**Supplemental Figure S1.** Flow-Cytometric DNA histograms for diploid and autotetraploid rice.

**Supplemental Figure S2.** Gene Ontology (GO) analysis of the up- and down-regulated genes in autotetraploid compared to diploid rice.

**Supplemental Figure S3.** Reproducibility analysis of Hi-C biological replicates.

**Supplemental Figure S4.** Interaction decay exponents of intra-chromosome arm interactions in each chromosome

**Supplemental Figure S5.** The TADs and loops are unrelated to transcription.

**Supplemental Figure S6.** The comparison of DNA methylation levels between 2C and 4C rice.

**Supplemental Figure S7.** DNA methylation levels in DEG promoters in 2C are similar to those in 4C rice, and CG and CHG methylation levels in non-DEG promoters in 2C rice are lower than those in 4C rice.

**Supplemental Figure S8.** The genomic distances between DEGs and differential methylation regions (DMRs).

**Supplemental Figure S9.** Average methylation level distribution over class I TEs.

**Supplemental Figure S10.** Average methylation level distribution over class II TEs.

**Supplemental Figure S11.** The methylation levels of TEs in promoters of DEGs and non-DEGs show no significant differences between 2C and 4C rice.

**Supplemental Figure S12.** Boxplots showing FPKMs between DEGs and non-DEGs in compartment A/B switched regions in 2C and 4C rice.

**Supplemental Figure S13.** The comparison of siRNAs between 2C and 4C rice.

**Supplemental Figure S14.** SiRNA abundance over TEs in TAD boundary genes and DEGs between 2C and 4C rice.

**Supplemental Figure S15.** The CG, CHG and CHH methylations in TEs of 2C and 4C rice.

**Supplemental Data Set S1.** List of DEGs in 4C rice compared to 2C rice.

**Supplemental Data Set S2.** GO-term categories of regulated genes in 4C rice compared to 2C rice.

**Supplemental Data Set S3.** The quality of Hi-C reads in 2C and 4C rice.

**Supplemental Data Set S4.** List of DEGs in chromatin compartment A/B switched regions.

**Supplemental Data Set S5.** List of non-DEGs in chromatin compartment A/B switched regions.

**Supplemental Data Set S6.** Bisulfite sequencing statistics.

**Supplemental Data Set S7.** List of TAD boundary genes in 2C and 4C rice.

**Supplemental Data Set S8.** The numbers of TAD boundary genes and DEGs within different distances to TEs or the numbers of TEs within different distances to TAD boundary genes and DEGs

## Acknowledgments

This work was supported by the National Key Research and Development Program of China (2016YFD0100902) and National Science Foundation of China (31871230 and 32170585 to Y.F. and 31971334 to Z.S.).

## Authors’ contributions

Z.S. and Y.F. designed the study; Z.S., Y.W., Z.S., H.Z., M.M., Z.T., P.W., YF., and D.C. performed the research; Z.S., Y.W., G.L., and Y.F. analyzed the data; Z.S. and Y.F. wrote the paper. All authors discussed the results and made comments on the manuscript.

